# Variant calling tool evaluation for variable size indel calling from next generation whole genome and targeted sequencing data

**DOI:** 10.1101/2021.07.15.452444

**Authors:** Ning Wang, Vladislav Lysenkov, Katri Orte, Veli Kairisto, Juhani Aakko, Sofia Khan, Laura L. Elo

**Author notes:** To whom correspondence should be addressed. Corresponding author: Laura L. Elo, Turku Bioscience Centre, Tykistökatu 6 A, FI-20520 Turku, Finland. Sofia Khan, Turku Bioscience Centre, Tykistökatu 6 A, FI-20520 Turku, Finland.

## Abstract

Insertions and deletions (indels) in human genomes are associated with a wide range of phenotypes, including various clinical disorders. High-throughput, next generation sequencing (NGS) technologies enable detection of short genetic variants, such as single nucleotide variants (SNVs) and indels. However, the variant calling accuracy for indels remains considerably lower than for SNVs. Here we present a comparative study of the performance of variant calling tools on indel calling, evaluated with a wide repertoire of NGS datasets. While there is no single optimal tool to suit all circumstances, our results demonstrate that the choice of variant calling tool greatly impacts the precision and recall of indel calling. Furthermore, to reliably detect indels, it is essential to choose NGS technologies that offer a long read length and high coverage, coupled with specific variant calling tools.

**Author summary:** The development of next generation sequencing (NGS) technologies and computational algorithms enabled large scale, simultaneous detection of wide range of genetic variants, such as single nucleotide variants as well as insertions and deletions (indels), which may confer potential clinical significance. Recently, many studies have been conducted to evaluate variant calling tools on indel calling. However, the optimal indel size range for different variant calling tools remain unclear. A good benchmarking dataset for indel calling evaluation should contain biologically representative high-confident indels with a wide size range and preferably come from various sequencing settings. In this article, we created a semi-simulated whole genome sequencing dataset where the sequencing data was computationally generated. The indels in the semi-simulated genome were incorporated from a real human sample to represent biologically realistic indels and to avoid inclusion of variants due to potential technical sequencing errors. Furthermore, we used three real-world NGS datasets generated by whole genome or targeted sequencing to further evaluate our candidate tools. Our results demonstrated that variant calling tools varies greatly in calling different sizes of indels. Deletion calling and insertion calling also showed differences among the tools. The sequencing settings in coverage and read length also had a great impact on indel calling. Our results suggest that the accurate indel calling was dependent on the combination of a variant calling tool, indel size range and sequencing settings.

## Introduction

Next-generation sequencing (NGS) has developed rapidly in recent decades. Compared with traditional Sanger capillary electrophoresis sequencing, or other directed polymerase chain reaction-based screening methods, it provides a more efficient and affordable way to detect genomic variants in large scale [1–3]. NGS is widely used in research and in the clinic [4–6] and consists of techniques to cover the whole genome, i.e. whole genome sequencing (WGS), or only exome regions, i.e. whole exome sequencing (WES), or certain genomic regions, i.e. targeted gene panel sequencing.

High-throughput NGS enables researchers to simultaneously identify large numbers of single nucleotide variants (SNVs) and insertions and deletions (indels), as well as other types of genomic aberrations, such as inversions and translocations. Indels are the second most common variant type in the human genome, after SNVs. They are also the most common type of structural variants (SVs), defined as genomic variants > 50bp [7, 8]. Indels have been implicated in many diseases, such as Parkinson’s disease and cancers, thus their detection in the human genome is significant for clinical research [4, 9].

To date, a number of computational tools for variant calling from NGS data have been published. They utilise different information such as concise idiosyncratic gapped alignment report (CIGAR) strings from binary alignment map (BAM) files and build their underlying algorithms based on paired-end sequencing reads, split-read, *de novo* sequence assembly, gapped sequence alignment or machine learning to detect indels. Paired-end reads methods, based on paired-end sequencing, use discordantly mapped paired-end reads to identify indel breakpoints, which are the junctions that define structurally variable genomic segments [10]. In essence, paired-end reads that are mapped further, or closer than the expected insert size in the library, may indicate that an indel occurred between the alignments of the two paired-end reads [10]. However, repeat regions in genomes, or SNVs near indel breakpoints, may influence the accuracy of indel calling results [10, 11]. Methods based on split-read take reads that span the breakpoint of an indel as evidence, to identify the indel at a single nucleotide-level [12]. However, the short read length produced by NGS lead to even shorter split reads, which limit the alignment and make it hard to obtain a sufficient read depth around the indel breakpoint to confidently call an indel [11]. Sequence assembly methods include *de novo* sequence assembly and local re-assembly. *De novo* sequence assembly assembles short reads into longer contigs, enabling fine-scale discovery of large indels, especially novel sequence insertions [13]. Local re-assembly takes reads around a potential variant site with reference sequence to re-build a haplotype and then generates an accurate variant allele and the corresponding genotype [14, 15]. However, *de novo* sequence assembly methods require more computational resources and are prone to assembly errors [11]. Gapped sequence alignment based methods use the alignment results from a gapped aligner, such as Burrows-Wheeler Aligner (BWA) [16], and apply probabilistic models to make indel calls [14, 17]. The mapping conditions of each base of reads from the input file provide evidence of indels. These methods require that the indels are contained within a read with correctly mapping conditions from the reads alignment step [18]. These methods are sufficient for detecting small indels which are fully covered by a single read length, but unsatisfactory for identifying indels longer than the read length [18]. Currently, as each of these methods have their own limitations, many variant calling tools use a combination of methods to detect a wider spectrum of indels [15, 19]. Lately machine learning methods such as the random forest model and the deep convolutional neural network have been applied in some of the variant calling tools to detect indels [20, 21].

In the past few years, many studies have been conducted to evaluate variant calling tools on indel calling, but these studies have only performed evaluations with limited selection of sequencing data types or only covered a limited indel size range. Sandmann et al. evaluated eight variant calling tools with both real and simulated non-matched targeted NGS data by evaluating the sensitivities, positive predictive values, F1 scores and other metrics of tools for calling SNVs and indels smaller than 50bp [22]. Supernat et al. compared three variant calling tools with WGS data of the well-known human individual NA12878, regarding variant calling precision, recall and F1 score in calling SNVs and short indels [23]. Zhao et al. evaluated three variant calling tools with real and simulated human germline WGS data by comparing precisions, recalls and F1 scores of candidate tools with different genome contexts [24]. However, these evaluation studies were only focusing on variant calling tools for calling small sized indels with a certain type of NGS data, the performance of tools for calling larger sized indels remains unknown. For large indel calling evaluation, Pei et al. used 14 next-generation and third-generation sequencing datasets to evaluate precision rates, recall rates and computation costs of 11 variant calling tools with both germline and somatic variant calling, but the effect of the varying indel size was not evaluated [25]. Kosugi et al. comprehensively evaluated 69 SV detection algorithms for different types of SVs by testing the representative tools with five simulated WGS datasets and a real WGS dataset for human individual NA12878 by using multiple evaluation metrics [26]. Cameron et al. selected 10 SV calling tools to evaluate their precision-recall rates, running times, concordances and quality scores by using three real WGS datasets and *in silico* datasets with different sequencing settings [27]. In those studies, comprehensive evaluations of variant calling tools were performed but focusing exclusively on SVs. The best performance of variant calling tools for certain size range of indels, for example, indels around 50bp still remains unclear. Each algorithm uses different evidence from sequencing data to detect the breakpoint of an indel. Therefore, the best detectable indel size range of algorithms might vary. It is important to determine the best detectable indel size range of different tools with different sequencing methods: a wrong selection may result in missed or false positive (FP) detections.

In this study, we performed an indel calling evaluation with variant calling tools that represent different types of methods. We tested them with a variety of sequencing data types, to discover the best indel calling size range and data type for each tool. This study uses both real sequencing data and simulated sequencing data. Simulated sequencing data can produce an accurate indel truth set, avoiding, as much as possible, any potential unmarked true positive (TP) indels in truth set which may cause erroneous false positive results from the detections of the tools. However, the production of simulated sequencing data, as *in silico* simulation processes, underestimates the potential technical errors in real world sample preparation and sequencing steps [28]. Real world sequencing data comes from real experiments and is indisputably the ideal data for tool evaluation. However, to our knowledge, there is no real sequencing data coupled with a truth set that covers a wide range of variable size indels, without technical bias, for example, tools involving truth set generation, that would be suitable for our evaluation. Therefore, this study used a semi-simulated WGS dataset with four different sequencing settings to evaluate eight candidate tools with a wide size range of indels which are adopted from the HuRef genome data [29]. In addition, we also used three real world sequencing datasets which are Genome in a Bottle (GIAB) NA24385 WES data [30], CHM1 cell line WGS data [31] and targeted gene panel sequencing data to evaluate the performance of tools on indel calling. Together with all the selected datasets, our study aimed to deliver an unbiased and comprehensive evaluation of variant calling tools on indel calling.

## Materials and methods

### Ethics statement

The targeted gene panel sequencing dataset in this study was obtained from leukaemias patients. The dataset was analyzed anonymously. The study involving human participants were reviewed and approved by the Ethics Committee of Turku University Hospital (approval no. 30/1802/2019) and Turku Clinical Research Centre (approval no. T012/014/19).

### Variant calling tools

For the evaluation, we selected eight widely used variant calling tools: DELLY [19], DeepVariant [21], FermiKit [32], GATK Haplotype Caller (GATK HC) [14], Pindel [12], Platypus [15], Strelka2 [20], and VarScan [17], which all represent different underlying methods for indel calling (Table 1). Below, the main features of each tool are described:

**Table 1.** Methodological strategies of the tools included in this study.

| Variants Calling tools | Paired-end reads | Split reads | Sequence assembly | Gapped sequence alignment | Machine learning |
| --- | --- | --- | --- | --- | --- |
| DeepVariant v0.7.0 |  |  |  | X | X |
| DELLY v0.7.9 | X | X |  |  |  |
| FermiKit v0.13 |  |  | X |  |  |
| GATK_HC v4.0.1.2<sup>a</sup> |  |  | X | X |  |
| Pindel v0.2.5.b9 |  | X |  |  |  |
| Platypus v0.8.1.1 |  |  | X | X |  |
| Strelka2 v2.9.2 |  |  |  | X | X |
| VarScan v2.4.3 |  |  |  | X |  |

<sup>a</sup>GATK\_HC: GATK Haplotypecaller

### DELLY

DELLY is a variant calling tool that is designed for SV discovery [19]. It uses aligned reads in a sorted, indexed, and duplicate-marked BAM file as an input. It outputs a binary variant call file, which can easily be converted to commonly used variant call format (VCF) [33, 34]. DELLY integrates paired-end reads method and split-read method, with data from different insert size libraries to call SVs at a single nucleotide resolution. DELLY can detect multiple types of SVs, such as deletions, tandem duplications, inversions, translocations and medium-size insertions, thus being able to call both large and medium-size indels.

### DeepVariant

DeepVariant is a variant calling software that makes use of a deep convolutional neural network [21]. It is implemented as an analysis pipeline which takes aligned reads in a BAM file as input with three steps: discovery and encoding of the candidate variants, genotyping using the neural network, and writing output in the VCF file. DeepVariant stacks so-called inception modules which concatenated multiple convolution filters. Upon processing, the output layer returns a probability distribution of genotype (homozygous and heterozygous) for each candidate variant. The non-reference sites are then assigned the most likely genotype and written into a VCF file. DeepVariant supplies two trained models for WGS and WES data predictions. The WGS model was trained on NA12878/HG001 and NA24631/HG005 samples, and the WES model was trained on NA12878/HG001 and NA24631/HG005 samples in version 0.7, respectively.

### FermiKit

FermiKit is a *de novo* sequence assembly based variant calling pipeline for Illumina whole-genome germline data [32]. It takes FASTQ files as input, assembles reads into contigs, and then maps them against a reference genome. It calls SNVs, short indels, and SVs from the reads alignment, with error correction. FermiKit uses several build-in modules to do error correction, *de novo* assembly, mapping and then parse the ‘pileup’ output and extract alignment break points to call SNVs, short indels and SVs in VCF format

### GATK Haplotype Caller

GATK HC is a variant calling tool that calls germline SNVs and indels via local re-assembly [14]. It first defines active regions from input BAM files that show differences with reference sequences, these active regions indicate the evidence of variants by using CIGAR information from BAM files such as mismatches, insertions or deletions in the mapped reads and the high base-quality soft clip reads. Then it splits reads from active regions into k-mers to identify candidate haplotypes by reassembling them with de-Bruijn-like graphs. After that, a pair Hidden Markov Model (pair-HMM) is built with state transition probabilities from the read base qualities to calculate the likelihood that each read was derived from each haplotype. The haplotype likelihood of each read from the pair-HMM is used to calculate raw genotype likelihoods using a Bayesian model. The genotype likelihoods are then used to call raw variants that are output in a VCF format.

### Pindel

Pindel is a split-read based pattern growth algorithm [35] that can detect the breakpoints of large deletions and medium-sized insertions from BAM files of paired-end short read sequencing data and report variants in VCF format with single nucleotide resolution [12]. Pindel first extracts the paired-end sequencing reads from the mapping results, keeping the paired reads that only one read can be mapped uniquely to the genome with no mismatch and the other read cannot be mapped to the reference genome under a certain alignment score. The 3 ′ end of the mapped read determines an anchor point. With the anchor point, a sub-region from the direction of the unmapped read will be searched with a user-defined maximum detection size. The 3′ and 5′ end fragments of the unmapped read will be used by the pattern growth algorithm to search possible unique substrings on reference genome within the certain size of a sub-region. The gap between 3 ′ and 5 ′ end fragments of the unmapped read is reported as an indel if at least two complete reads can be assembled with possible substrings from both 3′ and 5′ end fragments of unmapped reads.

### Platypus

Platypus is a haplotype based variant caller with local sequence assembly in a Bayesian statistical framework [15]. It uses BAM files as input, and outputs VCF files. First, it constructs candidate variants from the CIGAR strings of the BAM files, local re-assembly and external sources (VCF files provided by the user). Candidate variants are then assigned with pre-defined priors at the generation stage, and candidate haplotypes are generated. The likelihood of each haplotype is calculated with a HMM, by aligning a read to the haplotype sequence. Then, an expectation-maximisation algorithm is applied to estimate the haplotype frequency, using a diploid genotype model. With the haplotype frequencies, the posterior supports for variants are calculated by comparing the likelihoods between all haplotypes, including haplotypes which do not include a particular variant. Variants whose posterior support exceeds a threshold and pass a number of pre-defined filters are called.

### Strelka2

Strelka2 is a variant calling method for small variants [20]. It takes a BAM file as input and output a VCF file, and it is designed for both germline and somatic sequencing applications. The germline mode workflow defines several parameters from sequencing data and applies an indel error model to estimate error rates of indels and SNVs. It defines active regions where likely to have variants and uses alignments of reads or local *de novo* sequence assembly to generate candidate haplotypes and alleles which are passed the filtering. After that, candidate variants are phased, and genotyped by read re-alignment with a probability model. In the final step, Strelka2 applies pre-trained supervised random forest models for SNVs and indels, which are trained on the labelled data of the Platinum Genomes project [36] to improve variant calling precision.

### VarScan

VarScan takes a single mpileup file (text pileup file from BAM files) as an input, and outputs a VCF file [17]. It first scores and sorts the BAM file, discards reads which were mapped ambiguously to multiple positions or with a low identity, as well as unmapped reads where the aligner failed to map anywhere in the genome. The uniquely mapped reads are used to detect variants and determine the total number of reads that support each unique variant. VarScan then filters each predicted variant by the overall coverage, number of supporting reads, p-value, variant allele frequency, base quality, and the number of strands that are observed in the predicted positions of the variants.

### Datasets

To comprehensively evaluate the variant calling tools, we used both simulated sequencing data and real sequencing data (Table 2). We created a semi-simulated dataset with four different sequencing settings extracting a wide size range of indels from a real human sample, with known indels available as truth sets. Additionally, we utilized three real datasets that represented different sequencing data types. The details of datasets are described below.

**Table 2.** The semi-simulated WGS sequencing and real sequencing datasets used in this study.

| Data | Read length | Coverage | Data content | Usage | Sequencing platform |
| --- | --- | --- | --- | --- | --- |
| Semi-simulated data | 100bp | 5× | WGS data (chr1) | All sizes of indels | Illumina HiSeq 2000 |
|  | 100bp | 30× | WGS data (chr1) | All sizes of indels | Illumina HiSeq 2000 |
|  | 100bp | 60× | WGS data (chr1) | All sizes of indels | Illumina HiSeq 2000 |
|  | 250bp | 30× | WGS data (chr1) | All sizes of indels | Illumina MiSeq v3 |
| GIAB NA24385 data | 126bp | 135× | WES data | Small indels ($< 50\text{bp}$) | Illumina HiSeq 2500 |
| CHM1 cell line data | 101bp | 41× | WGS data | Large indels ($\geq 50\text{bp}$) | Illumina HiSeq PE-101 |
| Targeted gene panel sequencing data | 150bp | NextSeq: 2298× & MiSeq: 6651× | Targeted sequencing data | Clinical indels (3bp - 52bp) | Miseq v3 or NextSeq 500 v2 |

The National Center for Biotechnology Information recommends variants larger than 50bp to be submitted to dbVar, a database of human genomic structural variation, while variants less than 50bp should be submitted to dbSNP, a database containing human SNVs, small indels, and other types of small variants [37, 38]. Based on this separation, small indel calls (< 50bp) and large indel calls (≥ 50bp) from the real sequencing data were evaluated separately.

### The Semi-simulated WGS dataset covering a wide size range of indels with varying coverages and read lengths

In order to simulate as realistic data as possible with precise and fair indels for benchmarking purposes, we utilized HuRef genome data [29]. The original HuRef dataset was generated by Sanger sequencing and the variants were detected by Celera Assembler [29]. The use of Sanger sequencing may potentially limit mapping issues caused by short sequencing reads and an independent variant calling method for our candidate tools makes this data very suitable here. HuRef data also allowed us to reproduce a realistic dataset that would capture the challenges of small indel and large indel calling in human genome. If the indels were to be inserted randomly across the genome it would underestimate the proportion of indels located in ambiguous regions, where the indels may be represented in different positions [39] (S1 and S2 Figs). Here, we reconstructed chromosome 1 of the HuRef genome, based on human reference genome hg19, by similar methods to [39, 40] and inserted indels of a real human individual into the corresponding position in reference genome (S1 File). In addition, we reconstructed chromosome 1 with two different haplotypes by randomly selecting variants from different size ranges and only inserting them into one of the haplotypes as heterozygous variants or into both haplotypes as homozygous variants. In total, 20 692 insertions and 22 478 deletions, ranging from 1bp to 6053bp were included (S1 Table).

We created simulated paired-end sequencing reads using the NGS read simulation tool ART [41], with three different coverages: 5×, 30× and 60×. The read length of simulated paired-end sequencing data was 100bp. In addition, we created another simulated paired-end sequencing dataset with 30× coverage and 250bp read length, to compare how read length would affect indel calling. For each dataset, the sequencing coverage was contributed by the two haplotypes with an approximate ratio of 1:1, making it representative of a naturally diploid sample (S1 File). The semi-simulated paired-end sequencing dataset was publicly available at https://github.com/elolab/semi-simulated_indel_dataset.

### Genome in a Bottle NA24385 WES dataset for small indels (< 50bp)

The GIAB NA24385 dataset was used to assess small indel calling of tools from the real sequencing data. The GIAB datasets are considered as a gold standard among variant call sets. The high confidence in their variants is achieved by integrating and curating several call sets produced using competing sequencing platforms and variant calling tools. The variant datasets consist of seven individuals, of which one individual from the trio of Ashkenazi Jews (NA24385/HG002) was selected for this study [30]. The corresponding sequencing data was provided in a variety of conditions and technologies; the NA24385 WES data from Oslo University Hospital was selected for use in this study. Sequencing was performed on the Illumina HiSeq 2500 instrument, with Agilent SureSelect Human All Exon V5 capture kit, and yielded 150bp paired-end reads. The raw 135× sequencing data for GIAB NA24385 was downloaded using the Sequence Read Archive (SRA) toolkit with accession SRX1453593 [42]. In total, 5436 indels were used to evaluate the tools in this study.

### CHM1 cell line WGS dataset for large indels (≥ 50 bp)

The SV dataset of the CHM1 cell line was used to assess large indel calling of tools from real sequencing data. The CHM1 cell line is a haploid human hydatidiform mole lacking allelic variation [31]. The SV dataset of the CHM1 cell line was produced by a single-molecule, real time sequencing technology at a 54× sequencing coverage, which generated 18580 indels (≥ 50bp) that used in our study. All the sequence reads were aligned to GRCh37 using a modified version of the PacBio long read aligner and generated local assemblies by Celera and Quiver. Next, the SVs of CHM1 cell line were characterised systematically by a custom computational pipeline. The NGS short reads sequencing data of CHM1 cell line was produced on an Illumina HiSeq 2500, with 41× sequencing coverage and 101bp read length of paired-end reads. The raw sequencing data of CHM1 cell line from Illumina platform was downloaded by using the SRA toolkit with the accession SRX652547 [31].

### Targeted gene panel sequencing dataset

A total of eight targeted NGS datasets from patients with different leukaemias were collected from Turku University Hospital (TYKS), Finland. A targeted amplicon-based panel, TruSight Myeloid Sequencing panel (Illumina, US), was used to detect variants in diagnostic samples. The myeloid gene panel targets 54 genes with 568 amplicons, ranging from 225bp to 275bp in length. The combined coverage for libraries was 141 kb. For library preparation, 50 ng of DNA per sample was used, and the Illumina protocol was followed. The samples were pooled in series of 8 to 24 to obtain sufficient sequencing depth and 2 x 150 cycle sequencing runs were performed on the Illumina platform (Miseq v3 chemistry or NextSeq 500 v2 mid output chemistry). To analyse the quality of the sequencing run, parameters and data output from each run were compared against specifications outlined by the manufacturer (Illumina): cluster density (1200 – 1400 K/mm2 and 170 – 220 K/mm2 for MiSeq and NextSeq, respectively); Q30 greater than 75 % in both systems; and the total number of reads passing the filter and the total data yield of the run were evaluated to approve the data of the run. The average coverage of the amplicons in the data generated with the NextSeq 500 platform was 22987×, and for the MiSeq 6651×. As a truth set we used the somatic indels in genes *CALR* and *CEBPA* that were determined for diagnostic purposes in the Laboratory of Molecular Hematology and Pathology, TYKS Laboratory Division, Turku, Finland. The 52bp deletions in *CALR* exon 9 were analysed with capillary electrophoresis after PCR amplification, as described by [43]. The *CEBPA* indels ranging from 3bp to 36bp were determined as part of the diagnostic testing in Labor für Leukämiediagnostik, Munich, as described by [44].

### Evaluation criteria

We evaluated variant calling tools using their default running parameters, with some exceptions (S1 File). For data pre-processing such as sequencing reads alignment, we built a fixed bioinformatics pipeline to deal with both semi-simulated and real sequencing data (S1 File).

For the evaluation of indel calling with the semi-simulated WGS dataset, we first filtered out all the SNVs from every tool’s output, using vcftools [34], and only kept the indel results for further evaluation. We considered a tool-detected indel call as a TP if: 1) the position-match: the indel position deviation of a tool-detected indel and a truth-indel was between ±10% of the truth-indel size (the upper limit for position deviation was 50 bp), and 2) the size-match: the size difference between a tool-detected indel and a truth-indel was < 25% of the size of the truth-indel, and 3) the genotype-match: the genotypes of a tool-detected indel and a truth-indel are consistent. The tool-detected indels that failed to give genotype information were considered as FPs. In our truth set, there were no multi allele variants; if a tool called an indel as a multi alleles variant, this record was split as multiple records for each allele.

For GIAB NA24385 WES datasets, we used hap.py (https://github.com/Illumina/hap.py) to assess the evaluation results of each tool (S1 File). Hap.py is a VCF file comparison tool for diploid samples, which can benchmark variant calling results made by a variant calling tool against a truth set [45].

For the CHM1 cell line WGS datasets, we defined a TP as a tool-detected, filter-passed indel which has at least 20% overlap with a truth-indel via BEDtools [46] (S1 File). Criteria was kept loose because previous studies [39, 47] have reported that the variant calling methodology for CHM1 cell line WGS datasets has its own bias, which leads to a low concordance between tool-detected indels and truth-indels (S1 File).

The performance of tools was evaluated by calculating precision rate, recall rate, and F1 score. For clinical targeted gene panel sequencing data, due to the limited number of indels validated by clinical methods, we manually checked the results of each tool and summarised the results.

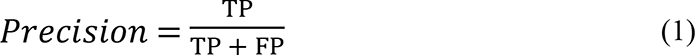

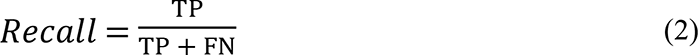

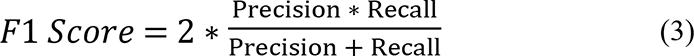

## Results

### Evaluation of variant calling tools of indel calling with various datasets

**Evaluation of small indel (< 50bp) and large indel (≥50bp) calling using the semi-simulated WGS dataset**

We evaluated the tool-detected indels of each tool for different sequencing coverages and read lengths with the semi-simulated dataset in five indel size ranges: 1bp – 20bp; 20bp – 50bp; 50bp – 200bp; 200bp – 500bp; > 500bp (Figs 1-6).

**Fig 1.**
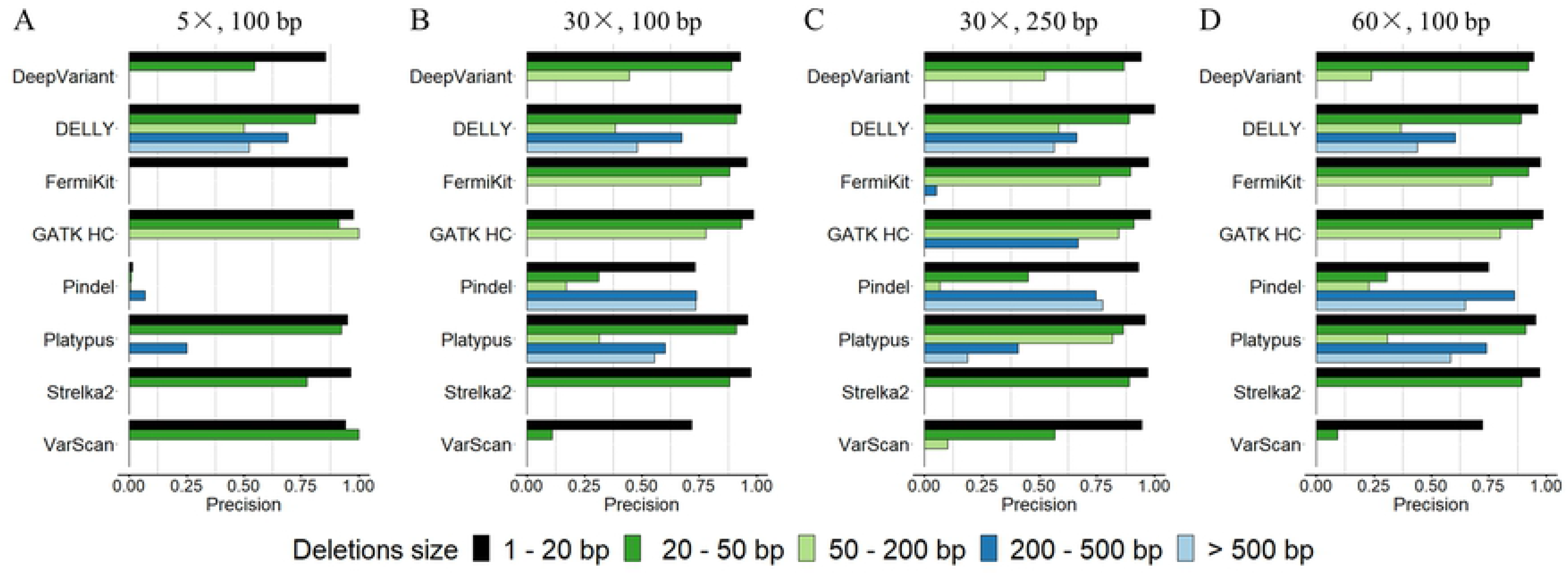
Precision rates of variant calling tools on deletion calling using the semi-simulated dataset. (A) 5× coverage, 100bp read length sequencing data. (B) 30× coverage, 100bp read length sequencing data. (C) 30× coverage, 250bp read length sequencing data. (D) 60× coverage, 100bp read length sequencing data.

Although with good precision rates in general, the majority of tools had relatively low recall rates with the 5× coverage sequencing data, especially FermiKit, and Pindel, whose recall rates of all indel size ranges were below 0.02 (Fig 1A, Fig 2A, Fig 4A and Fig 5A). With increasing sequencing coverage, the recall rates and the F1 scores of tools were improved, and the improvement from 5× to 30× was more obvious than the improvement from 30× to 60× (Figs 2-3 and Figs 5-6). Some very low recall rates made precision rates fluctuate greatly, but overall, the precision rates of tools showed less differences than the recall rates between coverages except for Pindel (Fig 1 and Fig 4). FermiKit and Pindel benefited the most with increasing coverages, which indicated that these two tools cannot work well with low coverage sequencing data. With the 5× coverage sequencing data, the F1 scores of DeepVariant, GATK HC, Platypus and Strelka2 for the indels in size range 1bp – 20bp were over 0.5, which indicated that for gapped sequence alignment based and machine learning based indel calling algorithms, the detections of indels smaller than 20bp might be more effective than indels larger than 20bp with low coverage sequencing data (Fig 3A and Fig 6A). DELLY also had F1 scores around 0.5 for deletions in size ranges 200bp – 500bp and > 500bp with the 5× coverage sequencing data, which showed that DELLY had relatively good abilities to call large deletion with low coverage sequencing data. In general, from the F1 scores, the results demonstrated that a higher sequencing coverage generates a better indel calling result (Fig 3 and Fig 6).

**Fig 2.**
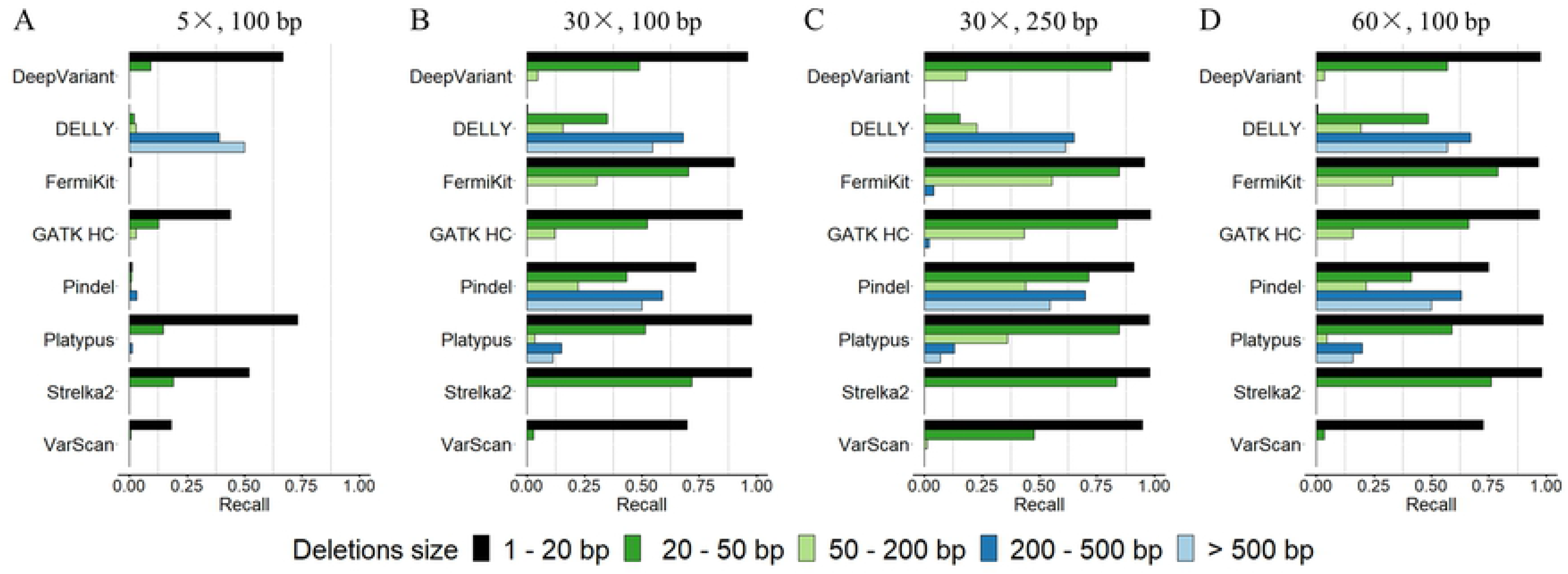
Recall rates of variant calling tools on deletion calling using the semi-simulated dataset. (A) 5× coverage, 100bp read length sequencing data. (B) 30× coverage, 100bp read length sequencing data. (C) 30× coverage, 250bp read length sequencing data. (D) 60× coverage, 100bp read length sequencing data.

**Fig 3.**
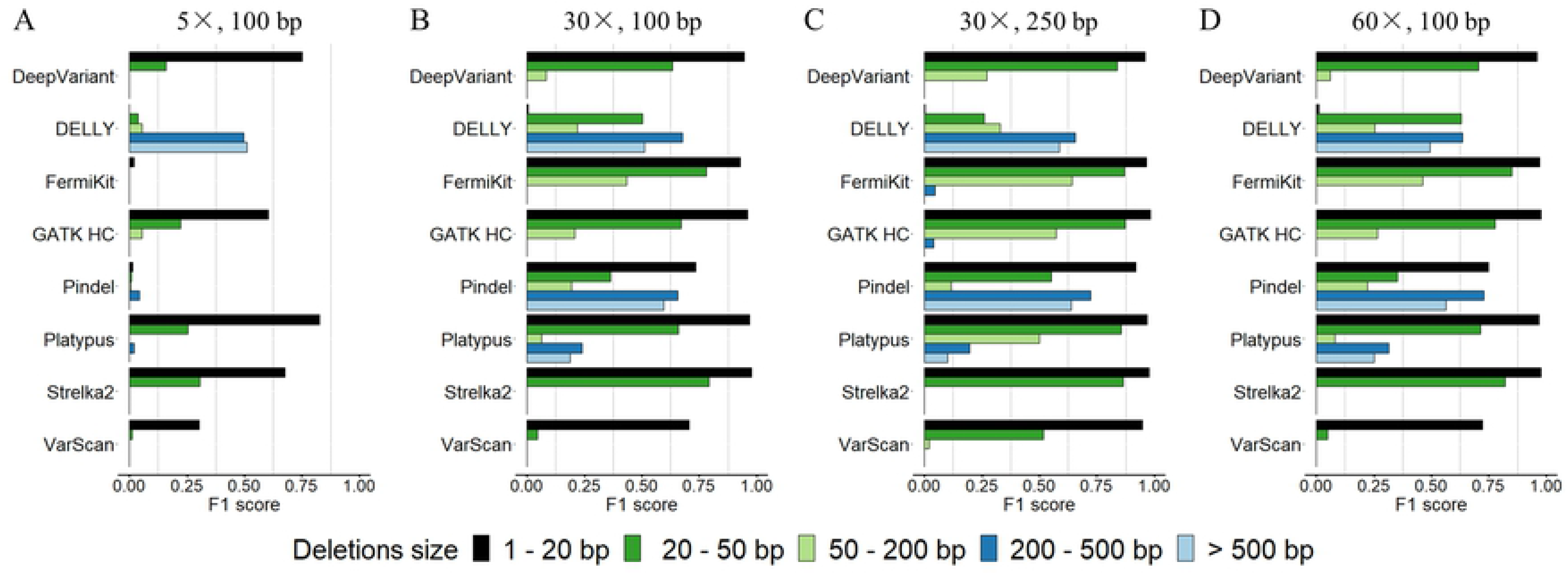
F1 scores of variant calling tools on deletion calling using the semi-simulated dataset. (A) 5× coverage, 100bp read length sequencing data. (B) 30× coverage, 100bp read length sequencing data. (C) 30× coverage, 250bp read length sequencing data. (D) 60× coverage, 100bp read length sequencing data.

**Fig 4.**
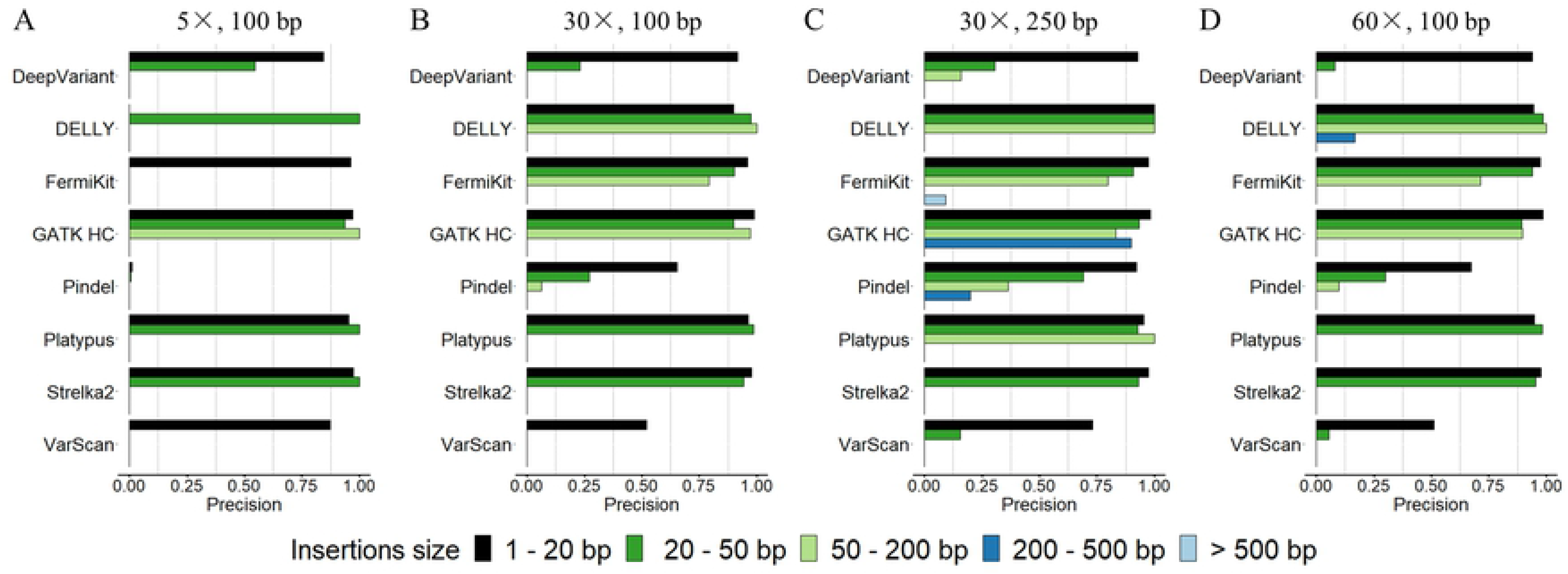
Precision rates of variant calling tools on insertion calling using the semi-simulated dataset. (A) 5× coverage, 100bp read length sequencing data. (B) 30× coverage, 100bp read length sequencing data. (C) 30× coverage, 250bp read length sequencing data. (D) 60× coverage, 100bp read length sequencing data.

**Fig 5.**
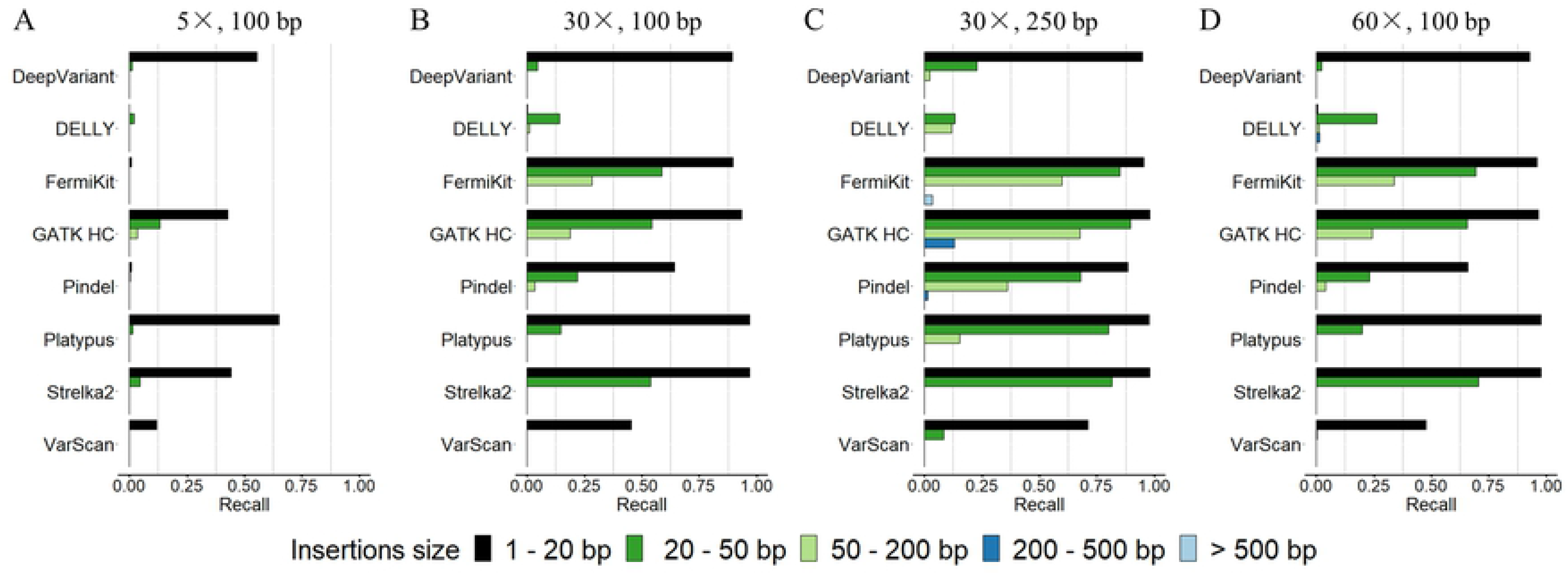
Recall rates of variant calling tools on insertion calling using the semi-simulated dataset. (A) 5× coverage, 100bp read length sequencing data. (B) 30× coverage, 100bp read length sequencing data. (C) 30× coverage, 250bp read length sequencing data. (D) 60× coverage, 100bp read length sequencing data.

For the assessment of how different read lengths affect indel calling, we evaluated the performance of tools by using the 100bp read length and the 250bp read length sequencing data with 30× coverage (Figs 1-6). We observed that the variant calling tools had wider detection ranges, better precision rates, recall rates and F1 scores with the 250bp read length sequencing data than with the 100bp read length sequencing data with the same coverage. The indel callings in size range > 50bp were remarkably improved. Both compared with 30× coverage 100bp read length sequencing data, the improvement of the precision rates, the recall rates and the F1 scores with 30× coverage 250bp read length sequencing data were slightly better than the improvement with 60× coverage, 100bp read length sequencing data. The results indicated that above a certain sequencing coverage, indel calling may benefit more by the increased the read length than the increased coverage of sequencing data. Although with low recall rates, DeepVariant, FermiKit, GATK HC, Platypus and VarScan detected larger indels that were not able to detect with 100bp read length sequencing data.

**Fig 6.**
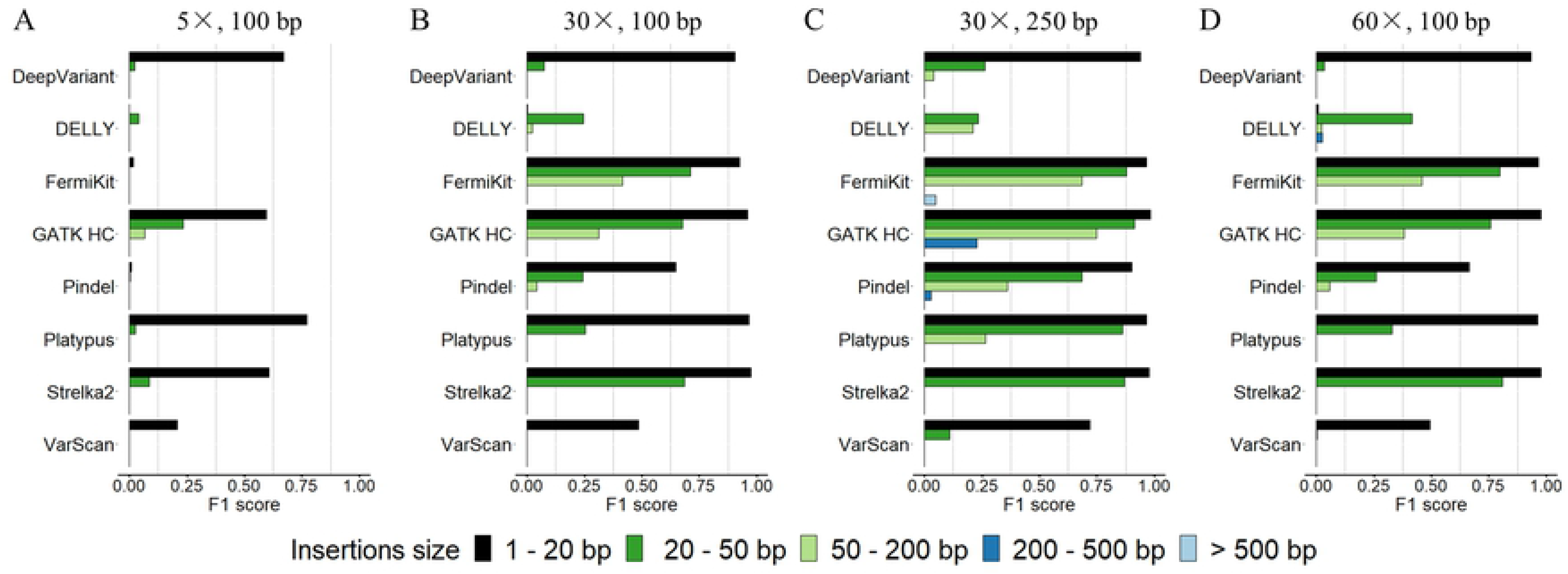
F1 scores of variant calling tools on insertion calling using the semi-simulated dataset. (A) 5× coverage, 100bp read length sequencing data. (B) 30× coverage, 100bp read length sequencing data. (C) 30× coverage, 250bp read length sequencing data. (D) 60× coverage, 100bp read length sequencing data.

From our results, we can see that tools with different algorithms showed different best variant calling size ranges. Taking the 30× coverage 100bp read length sequencing data as an example. For deletion calling with the 30× coverage 100bp read length sequencing data (Figs 1-3), for deletions in size range 1bp – 20bp, DeepVariant, FermiKit, GATK HC, Platypus and Strelka2 had both precision rates and recall rates over 0.9. The precision rates and recall rates of Pindel and VarScan for deletions in size range 1bp – 20bp were around 0.7. DELLY only detected limited number of deletions in size range 1bp – 20bp. For deletions in size range 20bp – 50bp, the recall rates decreased remarkably for the majority tools while DELLY started to detect deletions in this size range but with low recall rate. FermiKit and Strelka2 were the only two tools that had recall rates over 0.7 in this deletion size range. Except for Pindel and VarScan whose precision rates were low, the other tools all had precision rates over 0.9 for deletions in this size range. For deletions in size range 50bp – 200bp, a decrease was seen on both precision rates and recall rates for all the tools. FermiKit had the best F1 score on this deletion size range being little over 0.4. Strelka2 and VarScan failed to detect any deletions longer than 50bp. DELLY, Platypus and Pindel were the only three tools that detected deletions in size range 200bp – 500bp and > 500bp, Pindel had the best precision rate and DELLY had the best recall rate. Platypus had relatively good precision rates to detect deletions in this size range but recall rates were relatively low. For insertion calling with the 30× coverage 100bp read length sequencing data (Figs 4-6), DeepVariant, FermiKit, GATK HC, Platypus and Strelka2 had precision rates and recall rates around 0.9 for insertions in size range 1bp – 20bp. Pindel and VarScan had relatively low precision rates and recall rates while DELLY had very limited abilities to detect insertions in this size range. For insertions in size range 20bp – 50bp, notable decrease of recall rates was observed. Only FermiKit, GATK HC and Strelka2 had recall rates over 0.5. DELLY and Platypus had high precision rates but low recall rates for detecting insertions in this size range while DeepVariant and Pindel had both low precision rates and recall rates. VarScan was the only tool that failed to detect any insertion in this size range. For insertions in size range 50bp – 200bp, all the tools performed poorly. Only FermiKit and GATK HC had recall rates over 0.3 and all the other tools either had very low recall rates or even failed to detect insertions in this size range. No tool could detect insertions > 200bp. In general, tools performed better on deletion calling than insertion calling.

Previous research [48] has defined 50bp as the limit for SVs and small variants. We observed, however, that within the 1bp – 50bp range, the indel calling results were not consistent (Figs 1-6). For most of the tools with sequencing data of 100bp read length, there were notable decreases of precision rates and recall rates among indels in the size ranges 1bp – 20bp, 20bp – 50bp and 50bp – 200bp, especially for Pindel and VarScan. Compared with higher sequencing coverages, the longer read length sequencing data provided better precision rates and recall rates for longer indels sizes.

To further evaluate indel callings without genotypes taken into account, we relaxed our evaluation criteria to include indels that passed both position-match and size-match but that may have failed genotype-match (S3-S8 Figs). Without considering genotype-match, the precision rates and the recall rates of the tools were improved, especially for FermiKit, Pindel and VarScan. Although FermiKit still failed with 5× coverage sequencing data, it was able to call indels > 200bp and became the best tool to call insertions > 200bp with other semi-simulated sequencing data. The precision rates and the recall rates of Pindel were improved with all settings of the semi-simulated dataset and Pindel became the tool that had most comprehensive indel calling size ranges with 5× coverage sequencing data, which indicated that Pindel detected many indels with correct positions and sizes but failed to detect indels with correct genotypes. To further validate this hypothesis, we calculated the homozygous precisions, the heterozygous precisions and the proportions of indels with correct position-match and size-match but non-valid genotype information of tools with the semi-simulated WGS dataset (Fig 7). The results showed that Pindel with 5× coverage sequencing data had the largest proportion of non-valid genotype indel calls and it decreased dramatically when the sequencing coverage was increased. DeepVariant and FermiKit also had some non-valid genotype indel calls. In general, among the semi-simulated WGS dataset, the homozygous precision rates were higher than heterozygous precision rates. DeepVariant, Delly and VarScan had notable increase of heterozygous precision rates with the increased sequencing coverage. Pindel and VarScan also had an increase in heterozygous precision rates with the increased sequencing read length.

**Fig 7.**
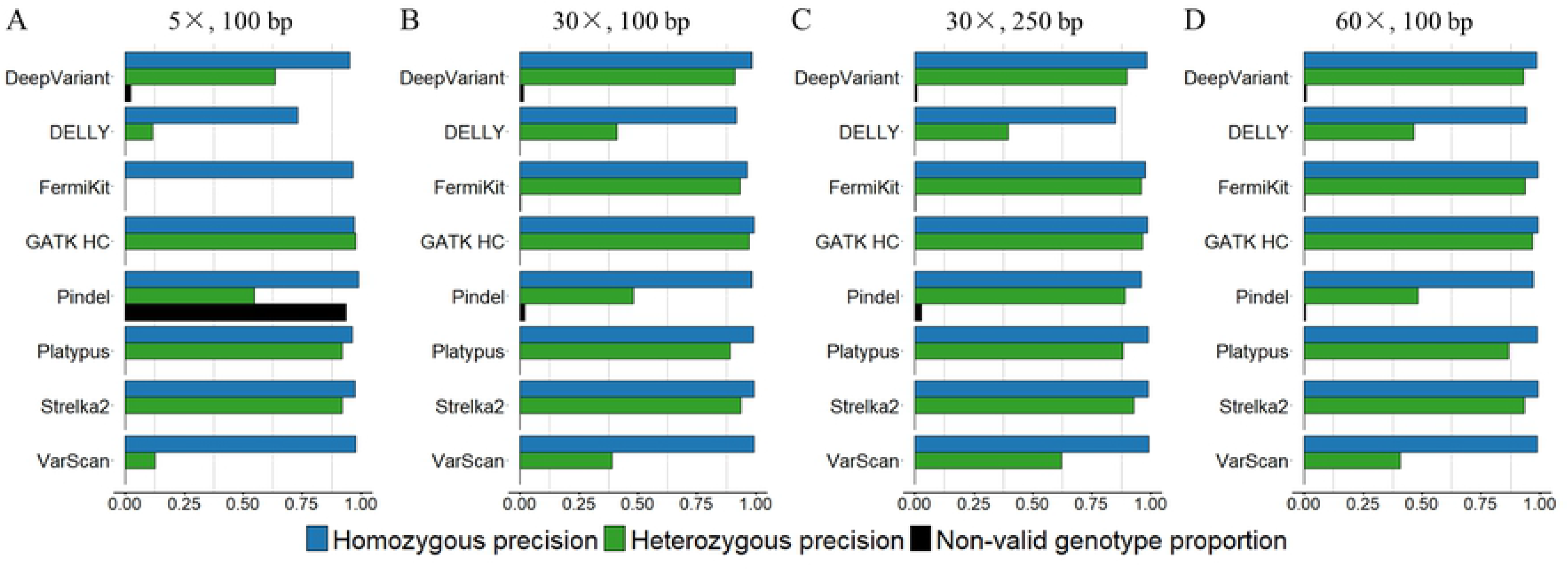
Homozygous precisions, heterozygous precisions, and non-valid genotypes proportions of variant calling tools using the semi-simulated dataset. (A) 5× coverage, 100bp read length sequencing data. (B) 30× coverage, 100bp read length sequencing data. (C) 30× coverage, 250bp read length sequencing data. (D) 60× coverage, 100bp read length sequencing data.

To better understand how the complexity of the genome sequence influences FP results, we gathered the FP results from the semi-simulated WGS dataset, and annotated FP indel calls based on their locations by using the “simple repeat” track from the UCSC genome browser via BEDtools. The results showed that indels located in simple repeat (SR) regions contains a large proportion of FP results, indicating that the breakpoint ambiguity caused by a SR region is the main challenge for indel calling (Fig 8). For 5× coverage sequencing data, the low read depth in indel breakpoints is the main reason that caused FPs. SR regions in genome may cause misalignment of reads and lead to further inaccurate indel calling, such as wrong positions or sizes due to the breakpoint ambiguity [39]. The breakpoint ambiguity of indels in SR regions may also cause incorrect allele frequency counting, thus leading to false genotype-level results (S9 Fig).

**Fig 8.**
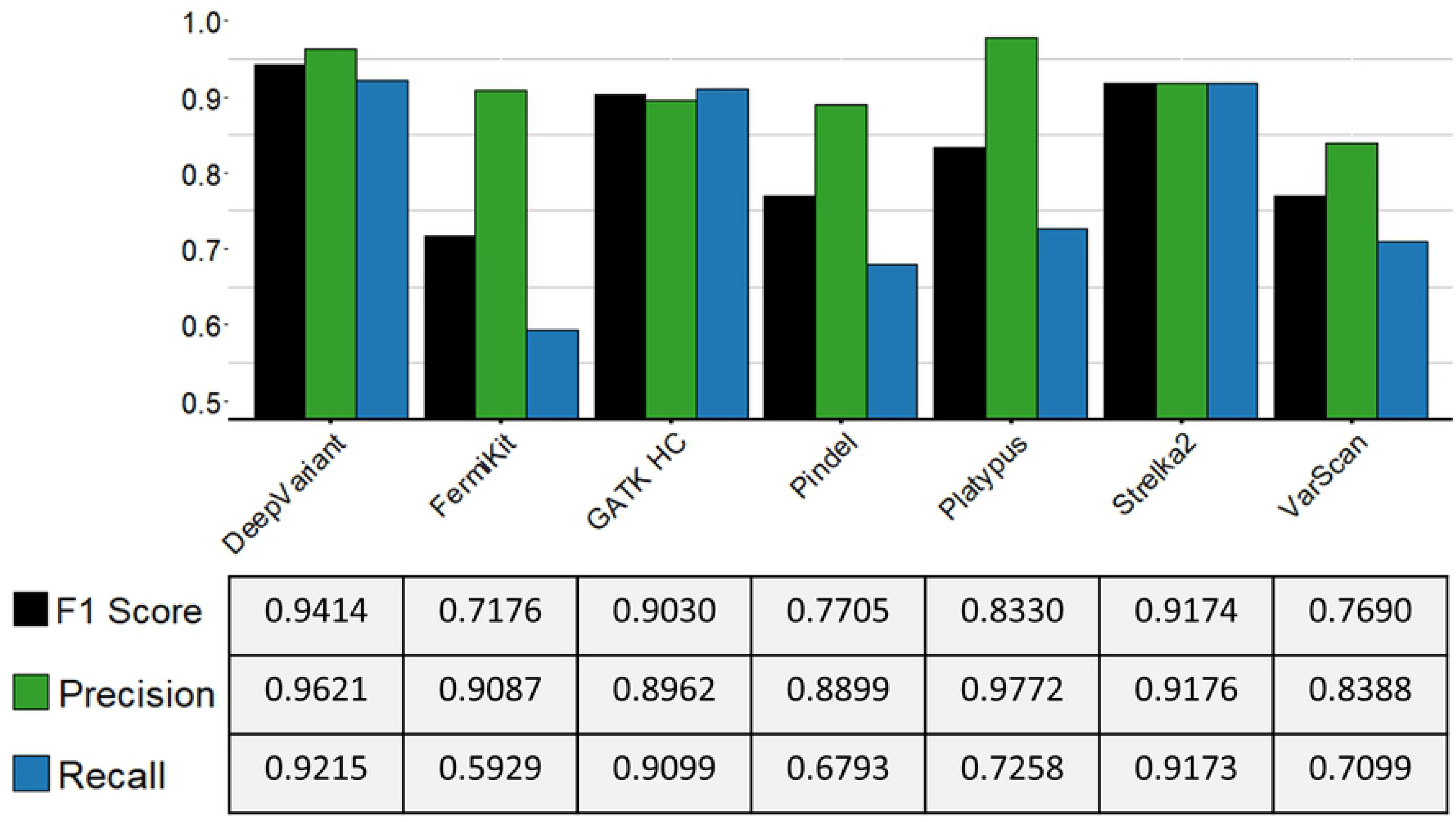
The proportions of SR and non-SR regions annotated FP indels of variant calling tools with the semi-simulated dataset. 5× coverage, 100bp read length sequencing data. (B) 30× coverage, 100bp read length sequencing data. (C) 30× coverage, 250bp read length sequencing data. (D) 60× coverage, 100bp read length sequencing data. The numbers on the top of each bar are the total numbers of FP results called by each tool with corresponding sequencing data.

### Evaluation of small indel calling using GIAB NA24385 WES data

From the F1 scores of tools with the semi-simulated WGS dataset, we selected the tools that were suitable for small indel calling (DeepVariant, FermiKit, GATK HC, Pindel, Platypus, Strelka2, and VarScan) for further evaluation with real sequencing data. The results in the GIAB NA24385 WES data (Fig 9) from hap.py showed that DeepVariant had the best recall rate and, together with Strelka2 and GATK HC, were the only three tools with recall rates > 0.9. Platypus had the best precision rate, followed by DeepVariant, both of which had precision rates > 0.95. DeepVariant also had the best F1 score, and along with Strelka2 and GATK HC, each had an F1 score > 0.9. At the genotype-level prediction (S2 Table), VarScan performed more poorly than the other tools. Pindel performed better genotyping with exome data than the semi-simulated WGS dataset, indicating that Pindel may perform better at genotype-level indel prediction with high coverage data. DeepVariant and Strelka2 had higher F1 scores than the other tools with the GIAB NA24385 WES data, which may indicate that machine learning tools have advantages of indel calling abilities over other indel callers.

**Fig 9.**
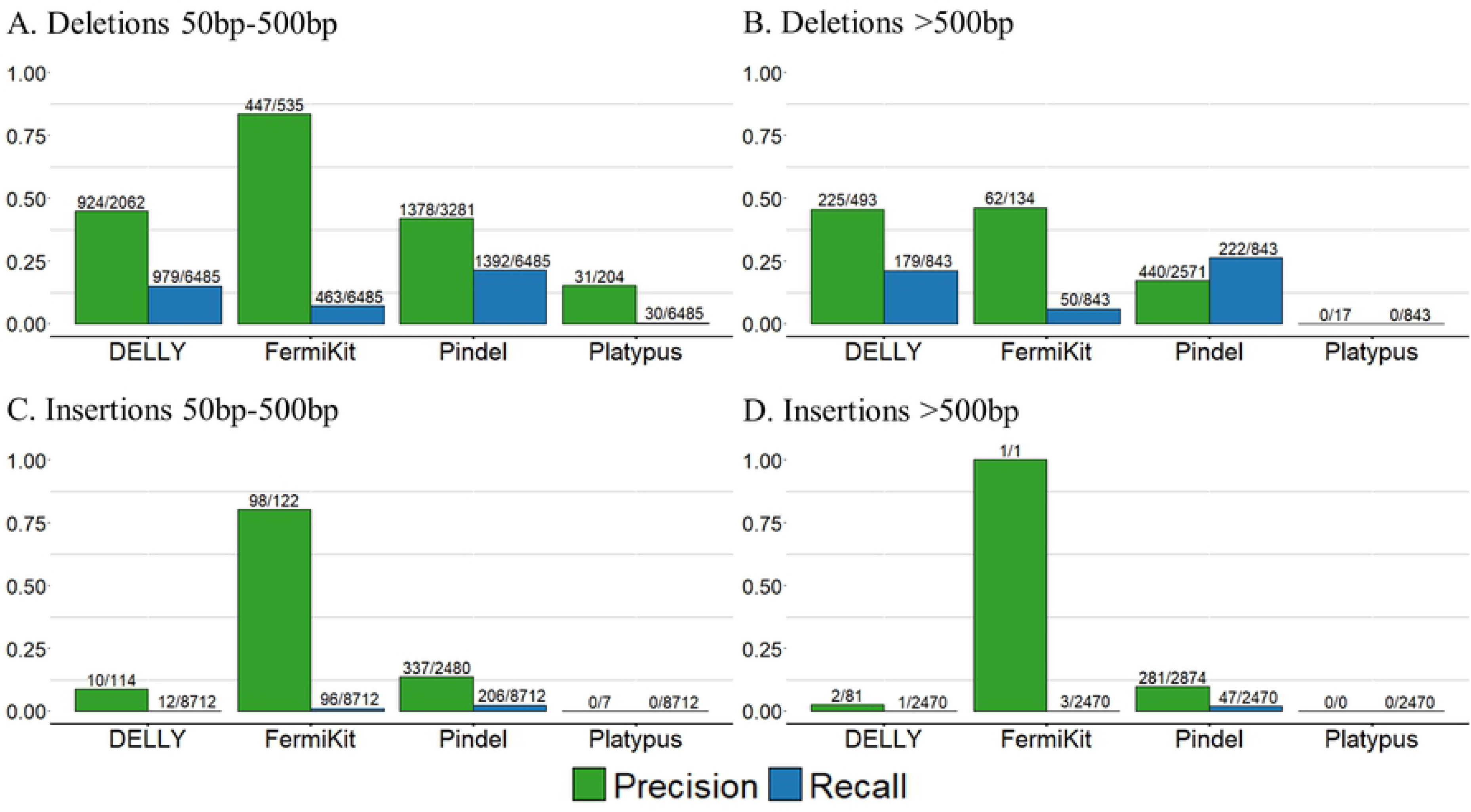
Indel calling evaluation results for variant calling tools with GIAB NA24385 WES data. The evaluation results were calculated using hap.py. The table below is the values of precision rates, recall rates and F1 scores of each tool.

### Evaluation of large indel calling using CHM1 cell line WGS data

From the results of the semi-simulated WGS dataset, we chose the tools that were suitable for large indel calling (DELLY, FermiKit, Platypus, and Pindel), and further evaluated them with real sequencing data. We used our pre-defined criteria to evaluate the performance of tools on large indels of size ranges 50bp-500bp, and >500bp in the CHM1 cell line WGS data (Fig 10). FermiKit had the best precision rates and Pindel had the best recall rates for indels of all size ranges. DELLY had comparable precision rates and recall rates for indels of all size ranges while Platypus had the worst performance and failed to call any insertion larger than 50bp. Although the overall variant calling of tools only called limited numbers of indels, the performance of tools on deletion calling was generally better than insertion calling and results with indels in size range 50bp-500bp were generally better than results with indels in size range >500bp Even though the criteria were loose, the concordance of the predicted results from the tools and the CHM1 truth set was still remarkably low. A possible reason for this is that variants in repeat regions can be called by CHM1 cell line PacBio calls, but cannot be detected efficiently by Illumina short-read sequencing [47]. A previous study which used CHM1 cell line dataset as a truth set, also suggested that the truth set might be missing variants [39].

**Fig 10.**
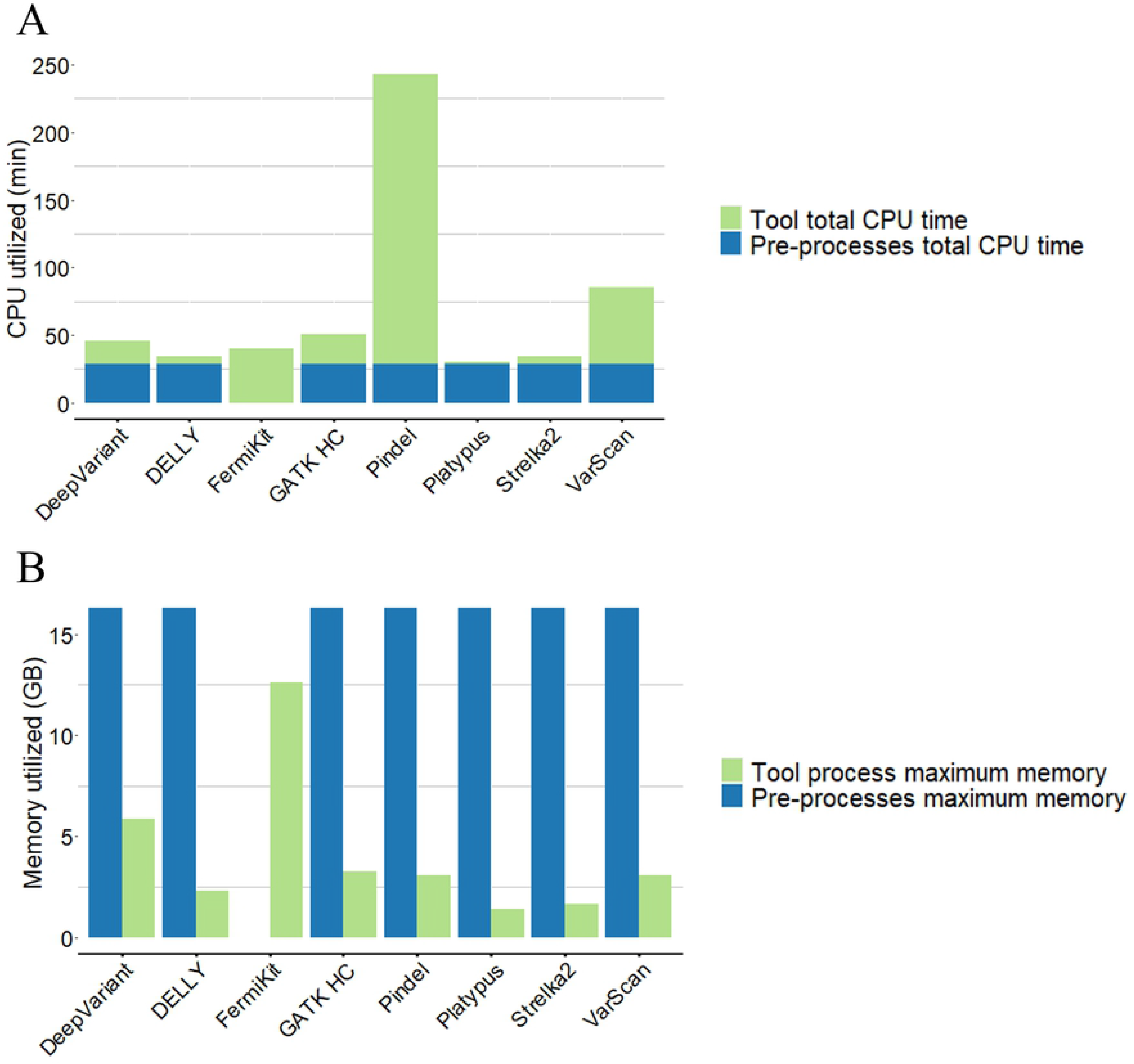
Indel calling results for variant calling tools with CHM1 cell line sequencing data. The results were separated into deletions (50bp – 500bp and > 500bp) and insertions (50bp – 500bp and > 500bp). The detailed precision rates and recall rates were shown on the top of each bar.

### Evaluation of indel calling using targeted gene panel sequencing data

We used amplicon-based clinical targeted gene panel sequencing data to further evaluate indels that ranged from 3bp to 52bp (Table 3). We only used the germline mode of variant calling tools and did indel calling in a non-matched variant calling manner due to the fact that we lacked the normal tissue sequencing data from the patients [49]. FermiKit is not designed for targeted sequencing data analysis, so it was not included in this evaluation. The results show that despite no tool detected the 22bp deletion of *CEBPA* gene from sample 1, DeepVariant, GATK HC, Pindel and Platypus successfully detected all the rest of the variants. DELLY and VarScan failed to detect two 3bp deletions of *CEBPA* gene and VarScan was the only tool that failed to detect one 36bp deletion of *CEBPA* gene. Strelka2 was the only tool failed to detect all four 52bp deletions of *CALR* gene.

**Table 3.** Indel calling results for the selected tools with targeted sequencing data of leukaemia patients.

| **Samples** | **Genes** | **Variants** | **DeepVariant** | **DELLY** | **GATK_HC** | **Pindel** | **Platypus** | **Strelka2** | **VarScan** |
| --- | --- | --- | --- | --- | --- | --- | --- | --- | --- |
| Sample 1 | *CEBPA* | 3bp insertion | Y | N | Y | Y | Y | Y | N |
|  | *CEBPA* | 22bp deletion | N | N | N | N | N | N | N |
| Sample 2 | *CALR* | 52bp deletion | Y | Y | Y | Y | Y | N | Y |
| Sample 3 | *CALR* | 52bp deletion | Y | Y | Y | Y | Y | N | Y |
| Sample 4 | *CALR* | 52bp deletion | Y | Y | Y | Y | Y | N | Y |
| Sample 5 | *CALR* | 52bp deletion | Y | Y | Y | Y | Y | N | Y |
| Sample 6 | *CEBPA* | 24bp insertion | Y | Y | Y | Y | Y | Y | Y |
| Sample 7 | *CEBPA* | 36bp deletion | Y | Y | Y | Y | Y | Y | N |
|  | *CEBPA* | 25bp deletion | Y | Y | Y | Y | Y | Y | Y |
| Sample 8 | *CEBPA* | 3bp insertion | Y | N | Y | Y | Y | Y | N |

For each sample there are one or more indel varying from 3bp to 52bp either in CEBPA or CALR. We manually evaluated the variant calling tools' results with our truth set. A successful call is marked with "Y", a failed call is marked with "N".

### Computational costs

We measured the running times and maximum memory usages of tools, based on one of the semi-simulated WGS data with 30× coverage and 250bp read length (Fig 11). All the indel calling processes, including read alignment and variant calling, were performed on a computer cluster managed by the free, open-source Simple Linux Utility for Resource Management. For each job we assigned 24 CPUs (Intel Xeon Processor, CPU MHz: 2095.078) per task, with 32G memory on the computer cluster.

**Fig 11.**
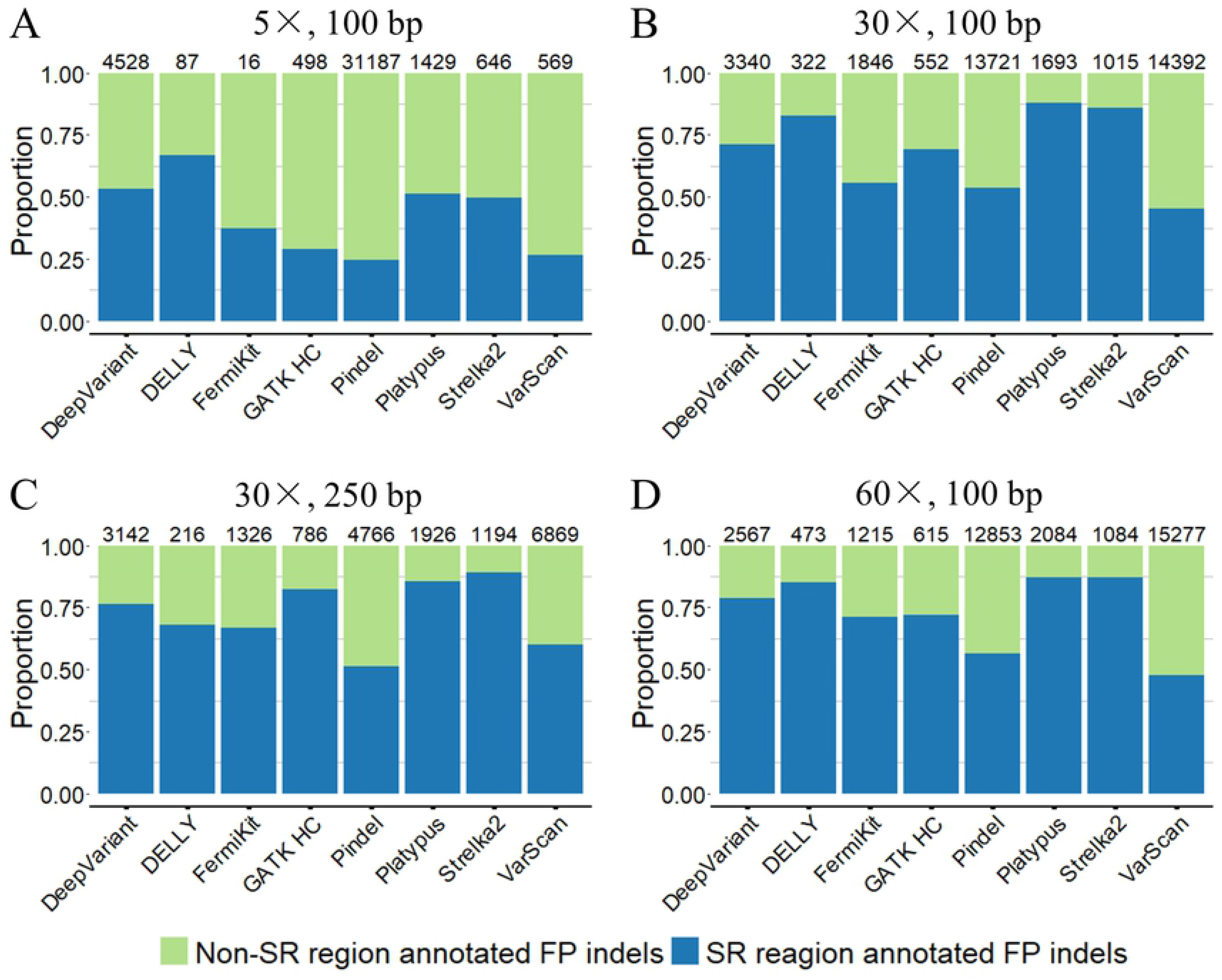
The running times and the maximum memory usages of variant calling tools. A) Total CPU times of each variant calling tool with 30× coverage, 250bp read length semi-simulated sequencing data. Pre-processes total CPU time included aligning the sequencing reads into the BAM file. Tool total CPU time included analysing the input BAM file into the output VCF format result. (B) Maximum memory usage of each variant calling tool with 30× coverage, 250bp read length semi-simulated sequencing data. Pre-processes maximum memory included aligning the sequencing reads into the BAM file. Tool maximum memory usage included analysing the input BAM file into the output VCF format result. Due to FermiKit is a *de novo* assembly algorithm based variant calling tool, which took sequencing reads as input and did not require pre-processes.

To save time and fully use high performance computational resources, parallelization is an important option for variant calling tools. Tools that are designed with a parallelization option are able to assign multiple CPUs to some steps of the data processing (DeepVariant, FermiKit, Pindel, Platypus and Strelka2). Tools without a parallelization option may only utilize one CPU to process the data (DELLY, GATK HC, and VarScan). From the results (Fig 11), Platypus was the fastest tool, while Pindel was the slowest. Platypus used the least memory, while FermiKit required the most. The high memory consumption of FermiKit is explained by reads alignment step that is included in the tool execution. The total utilized memory of FermiKit was lower than the memory consumption of the data pre-processes which was producing the sorted and indexed BAM files as the input for the other tools.

## Discussion

In this study, we evaluated eight variant calling tools for indel calling of different indel size ranges from different types NGS data. The tools represented different underlying algorithms including paired-end reads, split-read, *de novo* sequence assembly, gapped sequence alignment and/or machine learning based methods. The tools were tested using four different datasets with varying read coverage, read length and indel size ranges and that were produced with either Illumina HiSeq 2000, MiSeq v3, HiSeq PE-101 or NextSeq 500 v2 sequencing platforms. The semi-simulated WGS dataset with three different coverages (5×, 30× and 60×) and two different read lengths (100bp and 250bp) included small indels and large indels from 1bp to 6053bp. Since to our knowledge there is no single unbiased real world benchmarking dataset that would cover both small and large indels, we chose to use three real world datasets that represented WGS, WES and targeted gene panel sequencing data that together covers small and large insertions and deletions.

Our results demonstrated that the indel calling results varies greatly between insertions and deletions depending on size of indels and properties of sequencing data. Deep convolutional neural network and random forest model based tools, DeepVariant and Strelka2, demonstrated superior performance with small indel calling. However, their optimum indel call size range was limited to < 50bp. Split-read and paired-end reads based algorithms (DELLY and Pindel) could detect the breakpoints of deletions efficiently from the mapping information of sequencing reads. Although the precision rates and recall rates of large deletion callings were relatively lower than small deletion callings, these tools were best for deletion calling with various types of sequencing data. The *de novo* sequence assembly-based tool, FermiKit, had the best performance for large insertion calling. Due to the read length limitation of NGS, *de novo* sequence assembly algorithm could still be the best algorithm to detect large novel insertions in human genomes with WGS data, even though the performance of large insertion calling was not as good as small insertion calling. The local re-assembly methods (GATK HC and Platypus) were good for genotype-level indel calling. With the re-assembling of reads around indels, the context of variants could be better discovered, thus achieving greater genotype-level accuracies.

We evaluated the indel calling abilities and running times of our candidate tools. In addition, we also dissected the source of the FP indel calls. From our results, it can be seen that the majority of the FP calls came from SR regions that may be due to ambiguous sequence patterns around SR regions. Putative problems include misalignment and inconsistent representation of variants in mapping and variant calling steps, respectively. Nonetheless, we still may have underestimated the influence of SR regions because the difficulties of sequencing SR regions for sequencing platforms are not easily to be reproduced in our semi-simulated WGS dataset. In real sequencing data, even though the difficulties of sequencing error-prone regions can be reproduced, it is still difficult for different callers to reach a consensus with short reads, thus these regions are generally excluded from the confident regions of high-confident variant calls, e.g. [50]. Furthermore, in our semi-simulated WGS dataset, we shifted all the indels in the truth set to the left-most positions before the indel calling processes, because we considered any post normalization step with tool-detected indels such as left-aligning were extra burden for users, in addition some tools may gain extra advantage thus make evaluation unfair for the other tools. Even though we tried to avoid any inconsistent representation of indels by left-aligning all the indels in our truth set, it still might be that we underestimated the complexity of human genome sequences, which would require further efforts to investigate the ambiguity of indel calling.

In our evaluation, the majority of the tools were implemented with default parameters and only minor changes were introduced, when necessary. It is possible that the tools would benefit from optimised parameters to obtain more accurate results [51], but default parameters are the general starting point for tool usage, and allow an equivalent assessment of the performance of the tools. Moreover, optimisation requires modification of all possible combinations of the parameters, which is time-consuming and impractical [52]. Datasets may require different combinations of parameters, which makes it even more of a challenge to attain the universal best combination. Experienced users may have prior knowledge of how to optimise the tools, based on their data properties to obtain the best performance. In this study, we considered a user would use a given tool with its default parameters.

In general, higher coverage led to better results. The recall rates improved from 5× to 30×, then to 60×, especially for large indels. Fang et al. [53] suggested that 60× is the suitable sequencing coverage for WGS data from the HiSeq platform, however, in our study, we discovered that with the improved algorithms of indel calling methods in recent years, 30× sequencing coverage is also suitable in indel calling based on the performance of the tools. Our study showed no large differences in indel calling between 30× and 60× sequencing coverage data. Longer read length sequencing data was shown to help to call larger indels more accurately. In addition, longer read length may also contribute to alignment quality enabling more precise indel breakpoint detections. In our evaluation it was shown that indel calling for small size indels < 50bp were improved when machine learning based methods were applied. Although the precision and recall were still not quite on the same level as for the SNVs, the indel calling concerning small indels < 50bp were assuring. Future methodological development may benefit from improved machine learning models and *de novo* assembly methods to call also large, ≥50bp, insertions and deletions with good performance.

## Acknowledgements

We thank Elizabeth Nyman for language correction.

## Data availability statement

The semi-simulated dataset underlying this article are available in a GitHub repository at https://github.com/elolab/semi-simulated_indel_dataset. The truth set variants of semi-simulated dataset were adopted from HuRef genome (ftp://ftp.jcvi.org/pub/data/huref/). The Genome in a Bottle NA24385 WES dataset is available on https://www.nist.gov/programs-projects/genome-bottle. The WES data was downloaded using the Sequence Read Archive (SRA) toolkit with accession SRX1453593. The CHM1 cell line WGS dataset is available on https://eichlerlab.gs.washington.edu/publications/chm1-structural-variation/. The WGS data was downloaded by using the SRA toolkit with the accession SRX652547. All the bash scripts, python scripts, and R scripts which were used to generate results of this article are available at https://github.com/elolab/semi-simulated_indel_dataset. Targeted gene panel sequencing data used in this article cannot be publicly accessed due to patient confidentiality regulations. A permission for access to the targeted gene panel sequencing data has to be requested from the Turku Clinical Research Centre.

## Supporting information

**S1 Fig. An example of ambiguous variant.**

A deletion of “CA” was mutated in a simple repeat region with repeated pattern “CA”. In the truth set, the deletion was represented with left-align manner, but in tool prediction, the deletion was represented with right-align manner. These inconsistent representations caused a same single variant reported with different positions, reference alleles and alternative alleles, further causing trouble for evaluation.

**S2 Fig. Distribution of 10bp deletions by their positional ambiguities.**

The red bars represent Venter deletions and the blue bars represent deletions that were inserted at random sites.

**S3 Fig. Precision rates of variant calling tools on deletions callings evaluated without genotype-match by using the semi-simulated dataset.**

(A) 5× coverage, 100bp read length sequencing data. (B) 30× coverage, 100bp read length sequencing data. (C) 30× coverage, 250bp read length sequencing data. (D) 60× coverage, 100bp read length sequencing data.

**S4 Fig. Recall rates of variant calling tools on deletions callings evaluated without genotype-match by using the semi-simulated dataset**.

(A) 5× coverage, 100bp read length sequencing data. (B) 30× coverage, 100bp read length sequencing data. (C) 30× coverage, 250bp read length sequencing data. (D) 60× coverage, 100bp read length sequencing data.

**S5 Fig. F1 scores of variant calling tools on deletions callings evaluated without genotype-match by using the semi-simulated dataset.**

(A) 5× coverage, 100bp read length sequencing data. (B) 30× coverage, 100bp read length sequencing data. (C) 30× coverage, 250bp read length sequencing data. (D) 60× coverage, 100bp read length sequencing data.

**S6 Fig. Precision rates of variant tools on insertions callings evaluated without genotype-match by using the semi-simulated dataset.**

(A) 5× coverage, 100bp read length sequencing data. (B) 30× coverage, 100bp read length sequencing data. (C) 30× coverage, 250bp read length sequencing data. (D) 60× coverage, 100bp read length sequencing data.

**S7 Fig. Recall rates of variant calling tools on insertions callings evaluated without genotype-match by using the semi-simulated dataset.**

(A) 5× coverage, 100bp read length sequencing data. (B) 30× coverage, 100bp read length sequencing data. (C) 30× coverage, 250bp read length sequencing data. (D) 60× coverage, 100bp read length sequencing data.

**S8 Fig. F1 scores of variant calling tools on insertions callings evaluated without genotype-match by using the semi-simulated dataset.**

(A) 5× coverage, 100bp read length sequencing data. (B) 30× coverage, 100bp read length sequencing data. (C) 30× coverage, 250bp read length sequencing data. (D) 60× coverage, 100bp read length sequencing data.

**S9 Fig. Example of false genotyping caused by ambiguous regions.**

A homozygous deletion “chr1:878906 CTTT –→ C” was overlapped with 22 repeated “T” from chr1:878907 –-chr1:878928. The read that fully covered the repeat region had a 3bp gap in the CIGAR section of the BAM file. The read in which only the head or tail overlapped with the repeat region preferred to shorten its head or tail to omit the gap, based on the alignment algorithm. Simply counting the numbers of alleles at this site may lead to a low allele frequency, which then causes the tool to make a mistake and call a homozygous deletion a heterozygous one.

**S1 Table. Indel size distribution of the semi-simulated WGS dataset.**

Deletions and insertions are shown separately with different size ranges. The values are total number of variants with the number of heterozygous variants in brackets.

**S2 Table. Evaluation results of variant calling tools with GIAB NA24385 WES data.**

Summary evaluation results of variant calling tools with GIAB NA24358 WES data. Evaluation results were generated by hap.py.

**S1 File. Supplementary methods descriptions.**

